# Early Target Prediction in Action Observation

**DOI:** 10.1101/2025.07.28.667200

**Authors:** Martina Fanghella, Fabio A. D’Asaro, Davide Quarona, Guido Barchiesi, Marco Rabuffetti, Maurizio Ferrarin, Corrado Sinigaglia

**Author notes:** Corresponding Author Corrado Sinigaglia, Department of Philosophy, Università degli Studi di Milano, Via Festa del Perdono 7, 20122 Milano, Italy., Department of Philosophy, Stanford University, 450 Jane Stanford Way Building 90, Stanford, CA 94305.

## Abstract

Previous research has established that observers can predict action targets through hand preshaping. However, two critical questions remain unexplored: how predictions adapt to the available kinematic information and evolve throughout the movement timeline. We address these fundamental gaps by combining kinematic analysis with machine-learning approaches that differentiate between motor and visual cues. Using motion capture technology, we recorded reach-to-grasp actions toward large and small objects and had participants predict target size from hand kinematics at varying time points. Our analysis revealed that prediction performance not only evolved with increasing kinematic information but, crucially, differed significantly between target size choices. To provide insight into the underlying processes, we developed a comparative framework using two distinct machine learning approaches: Support Vector Machines (SVM) modeling kinematic information and CNN-RNN networks extracting visual patterns. The stronger alignment between human performance and SVM predictions offers empirical evidence that kinematic cues, rather than visual patterns, mostly guide target prediction. These findings advance our understanding of action prediction and have significant implications for social cognition and human-machine interaction.

**PUBLIC SIGNIFICANCE STATEMENT:** Understanding others’ intentions by observing their hand movements is crucial for social interaction, from passing objects to coordinating complex tasks. This study reveals that people use different cues to predict whether someone is reaching for a large versus a small object from the earliest stages of hand movement. By comparing human performance with Artificial Intelligence models, we found that people primarily rely on motor cues to make these predictions. These insights could improve rehabilitation techniques for individuals with social interaction difficulties and enhance the design of intuitive robotic assistants.

## INTRODUCTION

Navigating social interactions relies on tracking action targets. Knowing to which targets others’ actions are directed enables us to grasp their thoughts, plans, and intentions. We often identify these targets, even when hidden, by reading out the kinematic features of others’ actions (Becchio et al., 2012; Krishnan-Barman et al., 2017).

Consider hand actions. When grabbing an object, an agent has to transform its hand shape and size into the corresponding grip (Jeannerod et al., 1995). This correspondence has usually been investigated by appealing to kinematic variables such as the maximum grip aperture (i.e., the maximum distance reached by the index finger and thumb), the peak velocity of the wrist, and so forth. There is evidence that hand grip preshaping occurs well before reaching the target object, with both maximum grip aperture and its timing functionally related to object size (Jeannerod 1981, 1984; Jacobson & Goodale, 1991; Paulignan et al., 1991; Paulignan et al., 1997; Castiello, 2005).

Prior research has shown that an observer can predict what it is to be grasped by an agent just by capitalizing on the kinematic information conveyed by the acting hand preshaping. Ambrosini et al. (2011) found that observers can reliably identify intended targets by watching hand preshaping movements, enabling them to gaze at these targets before the hand arrives. They recorded saccadic movements while participants viewed an actor reaching for and grasping one of two objects of different sizes. In a control condition, the actor merely reached for and touched the objects holding their fist closed. In both conditions, participants had to look at the actions unfolding and had no prior knowledge about their potential targets. They turned out to be more accurate and faster in proactively gazing at the object to be grasped when they viewed the hand preshaping the grip according to the target size than when they viewed the hand reaching for the object with the closed fist (see also Ambrosini et al., 2012; Costantini et al., 2012a; and Costantini et al., 2012b).

A similar impact of hand preshaping on target detection has also been reported in early infants. 6-, 8- and 10-month-olds were tested while observing an agent grasp or touch a large or a small object (Ambrosini et al., 2013). All infants proactively gazed at the large target object when the observed hand preshaped a whole-hand grip. In contrast, only 8- and 10-month-olds proactively gazed at the small target with the observed hand preshaping a precision grip. Strikingly, this difference in proactive gazing correlated with the difference in infants’ ability to grasp large and small objects, preshaping the corresponding grip.

Ambrosini et al. (2015) expanded upon these findings by demonstrating that hand preshaping influenced both proactive gaze and explicit judgments regarding potential action targets, even in the presence of conflicting sources of evidence, such as the agent’s gaze looking in the opposite direction of the target. While participants initially tended to prioritize the agent’s gaze direction during early phases of action observation, the impact of hand preshaping became increasingly prominent as the observed action progressed, irrespective of the agent’s gaze direction.

Even more interestingly, Ansuini et al. (2016) showed that hand preshaping was critical in detecting potential action targets during the earliest phases of action observation. Unlike previous studies, they investigated the effect of target size over time by comparing the kinematic features of grasping movements toward small and large objects and the observers’ ability to identify the objects’ size from observing those movements. The grip aperture predicted the object sizes at 10% and 20% of the grasping movement time (Ansuini et al., 2015).

While kinematic features enable observers to identify action targets during grasping movements, fundamental questions remain about the underlying predictive process. How does prediction accuracy vary across different target types, and how does predictive strategy change throughout the movement timeline? We addressed these questions by systematically investigating how people predict hand action targets by tracking hand shape evolution over time, and how prediction performance relies on different cues for different targets at different timepoints. This approach moved beyond simple comparisons between hand kinematics and target identification to reveal the temporal dynamics and cognitive mechanisms underlying action prediction.

In doing this, we initially used motion capture techniques to record the kinematics of reaching for and grasping actions directed at large and small targets. Subsequently, we generated video clips depicting these actions as performed by a schematic upper limb by connecting the motion caption markers with red sticks. The participants’ task was to predict whether the observed hands grasped a large or a small object. Since no target object was visible, participants relied solely on hand shape changes over time to make their predictions. Moreover, the clips were shown up to four different time points to evaluate the information processing required for target prediction.

Our main aim was to investigate the prediction strategy employed to infer targets of different sizes over time, specifically exploring whether this strategy varied in the initial stages of target prediction, with such variations potentially linked to differences in relevant information cues. To address these questions, we implemented a three-tiered analytical approach. First, we quantified participants’ predictive performance through temporal analysis of detection sensitivity and target-specific accuracy metrics. Second, we developed a decision tree model to identify relationships between specific kinematic parameters and participants’ predictions across the movement timeline. Third, we introduced a novel comparative framework using two distinct machine-learning approaches to analyze how the target prediction strategy develops over time and explore the underlying information processing mechanisms and their key cues. We trained two distinct models: an SVM algorithm previously validated for motor control optimization and grip-based classification (Khera & Kumar, 2020; Ansuini et al., 2015), which used kinematic information as input, and a CNN-RNN architecture that extracted visual patterns through integrated spatiotemporal analysis directly from target videos (Karpathy et al., 2015). After evaluating target prediction performance for both models, we measured the agreement between the algorithms’ and participants’ performances to identify the key cues and processes driving target prediction, accounting for potential differences among targets, particularly in the early action observation phase.

## METHOD

### Transparency and Openness

Sensitivity analysis was computed with MorePower 6.0.4. (Campbell & Thompson, 2012). Statistical data analysis and Decision Tree were analyzed using SPSS, version 29.0.1.0. (IBM Corp. Released 2022. IBM SPSS Statistics for Windows, Armonk, NY: IBM Corp). SVM and CNN-RNN were implemented in Python, version 3 (Python Software Foundation, https://www.python.org/). Python packages are cited in the text and listed in the references section. Figures were created using R, version 4.1.2. (R Core Team, 2021, https://www.r-project.org/) and the package ggplot2, version 3.4.4 (Wickham, 2016, https://ggplot2.tidyverse.org/) and Excel (Microsoft Corporation, 2018. Microsoft Excel, https://office.microsoft.com/excel).

All data, analysis code, and research materials are available at https://osf.io/bdceu/files/osfstorage.

We report how we determined our sample size, all data exclusions, all manipulations, and all measures in the study.

The study’s design and its analysis were not pre-registered.

### Action Execution

Kinematic recording occurred at the LAMoBiR (“Laboratorio di Analisi del Movimento e Bioingegneria della Riabilitazione”), IRCCS Fondazione Don Carlo Gnocchi. Two agents (a female and a male) were asked to reach for and grasp, with their right (dominant) hand, two objects that differed in size (e.g., a large ball and a small ball, with diameters of 2 and 10 cm, respectively). At the beginning of each recording, the right hand was closed in a pinch and positioned 60 or 70 cm from the middle of the distance between the two objects (i.e., 10 cm). The two objects were randomly located on the left or right side, with the sides being counterbalanced for each target size. One hundred sixty actions were recorded (eighty for each target). A sound signal informed the agent to start the movement. Retro-reflective markers were positioned on anatomical landmarks of their right hand and arm according to the same procedure described in Carpinella et al., 2006. The 3-D spatial coordinates of the retro-reflective markers were captured with an optoelectronic SMART system (B|T|S, Milan, Italy) and then analyzed using a custom Matlab program (The Mathworks, Natick, MA, USA), as in Piedimonte et al. (2015).

### Target Prediction

#### Participants

Thirty-seven participants (mean age 22.68±33.68; 25 females) were enrolled in the present study. We initially set 40 as a required sample size, sufficient to detect a medium effect size (pη2= 0.088) given a power of .80 on the 2*4 repeated-measure ANOVA interaction of interest (Size*Time). Note that, in Ansuini et al., 2016, the effect of interest (2*4, Viewpoint*Time) was large (pη2= 0.24). Three participants had technical issues during the online testing session and, therefore, were excluded from the final sample. The final sample was composed of 37 participants, sufficient to detect a medium effect size (pη^2^= 0.095) on the interaction of interest, given a power of .80. All the participants were right-handed, had normal or corrected-to-normal vision, and had no history of psychiatric or neurological disorders. The Local Ethics Committee approved all research methods, which were carried out following the principles of the revised Helsinki Declaration (World Medical Association General Assembly, 2008). Written informed consent was obtained from all the participants.

#### Stimuli

The motion-capture recordings were edited to create one hundred sixty video clips (eighty for each target size). Red sticks connecting the motion-capture markers presented a schematic upper limb and hand. The target objects were not given. The video clips were acquired at 40 Hz and lasted 2230 ms on average. The presentation plane was the functional plane where the grip movement occurs.

We randomly selected a subset of sixteen video clips for the target prediction task, with a balanced representation of eight clips per action target. The selected clips differed in agent and target side to preserve action variability. All the chosen clips presented the targets at 70 cm. Frames from an example video clip are illustrated in Figure 1A (for two example video clips see also the Supplemental Material). The sixteen selected video clips were edited to represent four normalized time intervals (10%, 20%, 30%, 40%) of the movement time. We selected these four intervals because previous studies using up to 80% of movement time showed that at 40%, there was a discrimination rate comparable to longer (from 50% onwards) portions of the video (Ansuini et al., 2016). The movement time was calculated by excluding the reaction time of the agent for each trial.

**Figure 1.**
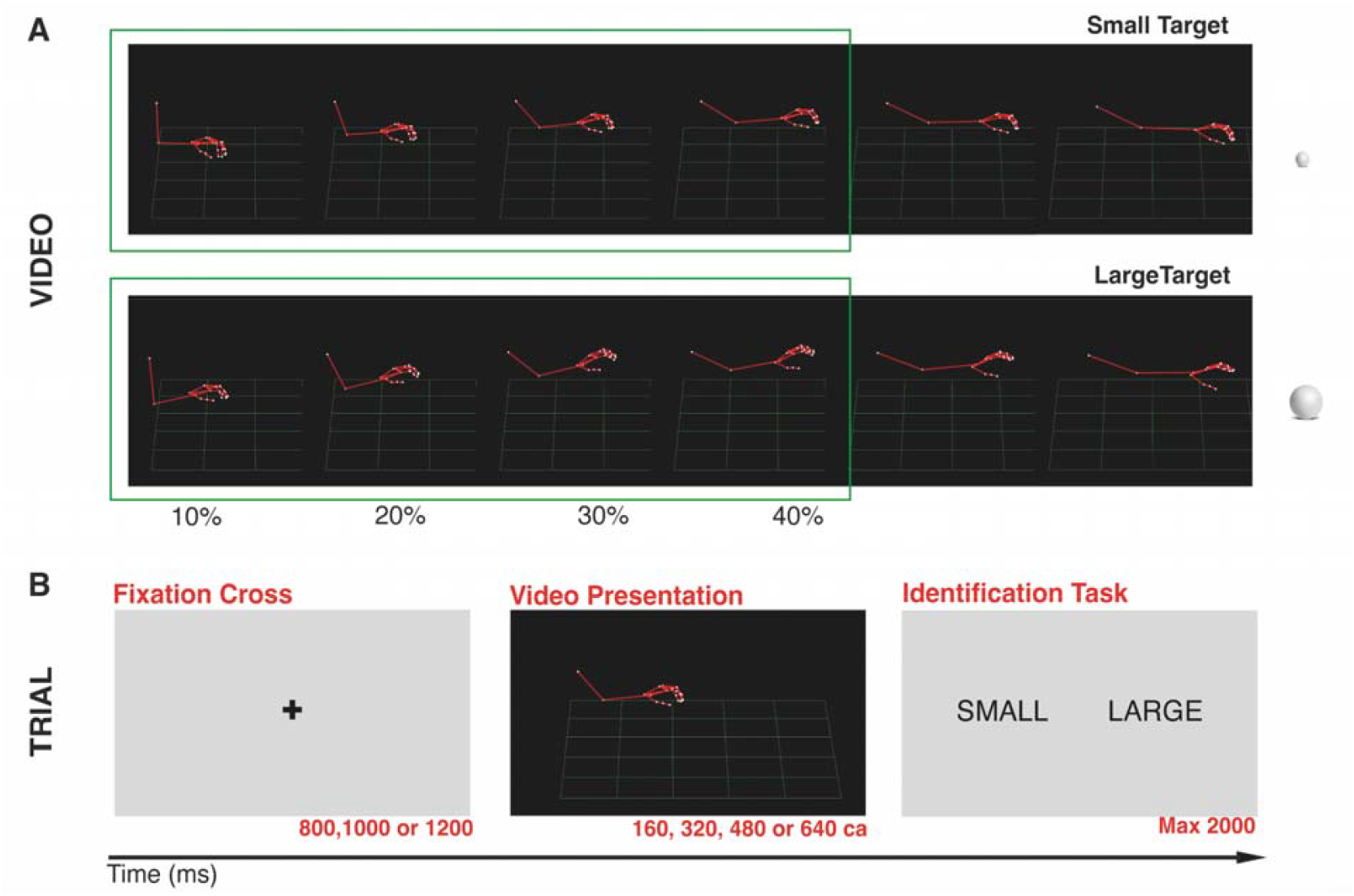
**A**. Example frames sampled from two acquired video streams at different times, one for the large ball and one for the small ball. Participants were shown up to 40% of these videos. **B**. Structure of a trial.

To rule out the possibility that the target sizes were merely mirrored by the duration of movements (Whitwell & Goodale, 2013), we compared grasping action durations for large and small targets for the selected video clips with a paired-sample t-test. Results showed that the grasping movements’ duration to large objects was not significantly different from that of the grasping movements directed to small objects (1564 ± 147 ms versus 1672 ± 122 ms; t(14) = -1.608, p = .130).

#### Procedure

Participants were asked to watch the selected 16 video clips and predict whether the observed hand grasped a large or a small object. Since the target objects were not presented in the video clips, participants could predict them by capitalizing on the preshaping of the grasping hands.

The trial started with a fixation cross of variable duration (800ms, 1000ms, and 1200ms), followed by the video clip presentation with various time intervals (respectively, the initial 10%, 20%, 30%, and 40% of the movement time), and a response slide (2000 ms) where participants had to decide if they observed a small or a large object grasping action, by stroking a “n” or a “m” key (response keys were counterbalanced across participants). For a graphical depiction of a trial, see Figure 1B. The experiment was run in 8 blocks of 64 trials (each video clip was repeated eight times for each of the four initial movement times in randomized order) and pause time between blocks lasted from 1 to 5 minutes.

The experiment was conducted online using participants’ computers and the E-prime Go software. At the beginning of the task, the experimenter contacted the participants via Microsoft Teams and provided oral and written instructions.

### Data Analysis

#### Action Execution

A custom Matlab script extracted the grip aperture, defined as the distance between the markers on the thumb and index tips during the reach-to-grasp actions (see Piedimonte et al., 2015), and wrist transport velocity. A 2×4 repeated-measure ANOVA was run on grip aperture and wrist velocity for the reach-to-grasp actions represented in the 16 video clips with the within factors Size (Large/Small) and Time (10%, 20%, 30%, 40% of the movement time). As a sanity check, we computed a 2×2×2×2×4 mixed repeated-measure ANOVA on grip aperture and wrist velocity on the whole dataset (160 executed actions) with the between-factor Agent (Male/Female) and the within factors Side (Left/Right), Distance (60/70), Size (Large/Small) and Time (10%, 20%, 30%, 40%).

#### Target Prediction

Participants’ responses and Reaction Times (RTs) were collected using E-prime Go software. Correction of 2.5 SD for RTs was applied at the participant level (i.e., the analysis excluded all trials with RTs above or below 2.5 SD from mean RTs). Following Ansuini et al., 2016, we calculated the *d’*, which measures target prediction performance (Green & Swets, 1966). After calculating *d’* for each participant, we computed a 4-level repeated-measure ANOVA with within-factor Time (10%, 20%, 30%, 40% of the movement time).

Then, we calculated two target-related performance metrics, Recall and Precision, using the standard formulas:

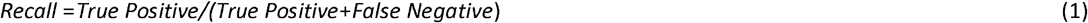

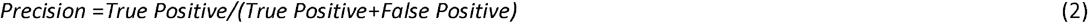

For example, the Recall of the Small response is calculated by considering the number of times Small is answered and the answer is true, divided by the sum of the number of times Small is answered and the answer is true and the number of times not Small is answered, and the not Small response is false. Meanwhile, the Precision for the Small response is calculated by considering the number of times Small is answered, and the answer is true, divided by the sum of the number of times the Small response is answered, and the answer is true, and the number of times Small is answered, and the answer is false. The same holds for the Large responses.

We conducted two 2×4 repeated-measure ANOVA on participants’ Recall and Precision, including the within-factors Size (Large/Small) and Time (10%, 20%, 30%, and 40% of the movement time). Finally, we tested differences in RTs with a 2×4 repeated-measure ANOVA, including the within-factors Size (Large/Small) and Time (10%, 20%, 30%, and 40% of the movement time).. Only RTs for correct responses were considered in the analysis.

When Sphericity was violated, we applied Greenhouse-Geisser correction. All post-hoc tests are Bonferroni-corrected for multiple comparisons.

#### Decision Tree

We analyzed participants’ responses with a Decision Tree (DT) using the SPSS software. This algorithm allowed us to predict participants’ performance based on the kinematic parameters recorded during action execution. Our dependent variables were participants’ responses (Large/Small) at 10%, 20%, 30%, and 40% of the movement time, and our predictors were grip aperture and wrist velocity.

#### Classification

We wanted to understand which cues and processes drive participants’ prediction strategies. We trained two algorithms to classify reach-to-grasp targets (Large/Small) and compared the Recall and Precision of their responses with participants’ performance. Both algorithms were trained on the dataset consisting of 70% of the 160 actions (i.e., 112) and tested on the remaining 30% (i.e., 48).

The first algorithm, Support Vector Machine (SVM), is a classical Machine Learning method usually applied to small-to-medium-sized datasets (Cristianini & Shawe-Taylor, 2000). This algorithm has previously demonstrated success in optimizing motor control through kinematic parameter analysis. For instance, SVM has been shown to be the best classifier for gait analysis, providing valuable insights for gait diagnosis and rehabilitation (Khera & Kumar, 2020). Furthermore, Ansuini et al. (2015) took advantage of SVM to show that kinematic parameters such as grip aperture and wrist velocity function as predictors of target size from the early stages of action execution. In the present study, SVM was trained and tested on the same kinematic parameters extracted from the execution of the reach-to-grasp actions. Training and tests were performed at each movement time (10%, 20%, 30%, and 40).

The second algorithm is a neural network with a CNN-RNN architecture, selected for its capacity to model spatial and temporal dependencies in video data. While this architecture is not biologically faithful, it draws conceptual inspiration from hierarchical processing in the visual system (Lindsay, 2021) and the role of temporal context in perception (Lipton, 2015). CNNs and RNNs together provide a practical framework for analyzing dynamic visual sequences, as evidenced by prior work demonstrating their utility in modeling aspects of perceptual input (Karpathy et al., 2015). It is worth noting that our aim is similar, e.g., to Bruera and Poesio (2024), who use large language models to predict brain activation patterns without claiming those models are mechanistically brain-like; our use of CNN-RNNs aims to capture task-relevant structure in the data, not to serve as a biologically precise account of human vision. We acknowledge the limitations of such analogies (Wichmann & Geirhos, 2023) and emphasize that our model is evaluated based on performance in a behavioral task, not its neural plausibility. The algorithm was trained and tested on video frames corresponding to four selected movement times.

We used both algorithms’ freely available Python 3 implementations, specifically, the svm package from the sklearn library (Pedregosa et al., 2011, available at: https://scikit-learn.org/stable/) and the VGG16 model from the keras library included in tensorflow (Kapoor et al, 2023; available at: https://keras.io/).

We experimented with different hyperparameter choices regarding the SVM model and found the RBF kernel to perform best. We left the other parameters to their default values (for reference, default parameters can be found at https://scikit-learn.org/stable/modules/generated/sklearn.svm.SVC.html).

For our CNN-RNN architecture, we implemented two GRU layers followed by a dropout layer for regularization and a dense output layer with sparse categorical cross-entropy loss. Hyperparameters were optimized with a grid search across the following ranges: layer units (8, 16, 32, or 64 for each layer), dropout rates (0.2 and 0.4), optimizers (Adam and RMSprop), activation functions (tanh, ReLU, and sigmoid for dense layers, with softmax for output), and learning rates (0.001 and 0.0001). The optimal configuration consisted of 32, 16, and 8 neurons in the first, second, and third layers respectively, with a 0.4 dropout rate, ReLU activation in the dense layer, softmax activation in the output layer, Adam optimizer, and 0.001 learning rate. In both cases, we then performed 10-fold cross-validation for evaluation.

#### Light Cohen’s Kappa

We quantified agreement between the SVM and CNN-RNN algorithms and participants with a metric known as *Cohen’s Kappa*, first introduced to measure agreement between observers of psychological behaviors (Cohen, 1960). Mathematically, Cohen’s Kappa is defined as 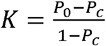, where *P*_0_ is the total probability of agreement between raters, and *P*_c_ is the probability of such agreement occurring due to chance. To illustrate, imagine that a human participant *H* and an SVM *C* classified the same 16 videos. Furthermore, assume that seven videos were classified as “Small” by both *H* and *C*, four videos were classified as “Large” by both *H* and *C*, two videos were classified as “Small” by *H* and “Large” by *C*, and the remaining three videos were classified as “Large” by *H* and “Small” by *C*. Then, 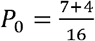 and 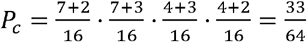, and 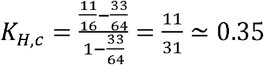, which is considered a “fair” agreement between *H* and *C*.

To match the number of participants, we trained each algorithm (SVM and CNN-RNN) 37 times on distinct random subsets of the training dataset. We got 37 different classifications of the 16 test reach-to-grasp actions at 10%, 20%, 30%, and 40% of the movement time. Since participants evaluated each test video clip eight times for each movement time, we considered the mode of their responses as their final evaluation of the video. For example, if a participant classified a video clip at 20% of the movement time five times as “Small” and three times as “Large,” we consider “Small” as its final classification. In the case of a tie, we arbitrarily assign this to class “Small” – this does not affect the results as ties are scarce.

We calculated Cohen’s Kappa and averaged the results to measure the agreement between the participants and the model for each pair of a trained model and a participant. Note that average Cohen’s Kappa is sometimes called *Light Cohen’s Kappa* (Light, 1971): we use this terminology below. When interpreting Cohen’s Kappa (including Light Cohen’s Kappa) values qualitatively, Fleiss et al. (2003) suggest that these values can be associated with intuitive meanings, as shown in Table 1. We will use this intuitive interpretation when presenting the results.

**Table 1.** Qualitative interpretation of (Light) Cohen’s Kappa according to Fleiss et al. (2003).

| Value of K | Qualitative interpretation |
| --- | --- |
| $< 0.4$ | Poor agreement |
| $0.40 \leq K < 0.6$ | Fair agreement |
| $0.6 \leq K < 0.75$ | Good agreement |
| $0.75 \leq K < 1$ | Excellent agreement |

## RESULTS

### Action Execution

#### Grip aperture

In the main 2×4 repeated-measure ANOVA, we found a main effect of Time (*F*(3,18)= 121.147, *p* < .001, *pη* = 0.953) and Size (*F*(1,6)= 79.584, p <.001, *pη* = 0.930), as well as a statistically significant Size*Time interaction (*F*(3,18)= 95.868, p < .001, *pη* = 0.941). Post-hoc test with Bonferroni correction on the Size*Time interaction revealed a greater grip aperture for the Large target than for the Small target from 20% up to 40% (mean±SE: 45.582±3.111 vs 30.30±1.06mm, p =.026 at 20%, 68.858±4.23 vs. 33.183±1.48mm, p<.001 at 30%; 84.791±4.60 vs. 35.281±1.78mm, p<.001 at 40%). We did not find differences in grip aperture between the Large and the Small target at 10% of the movement time (28.786±1.67mm vs 26.752±0.48mm, p = 1.). Results are shown in Figure 2A.

**Figure 2.**
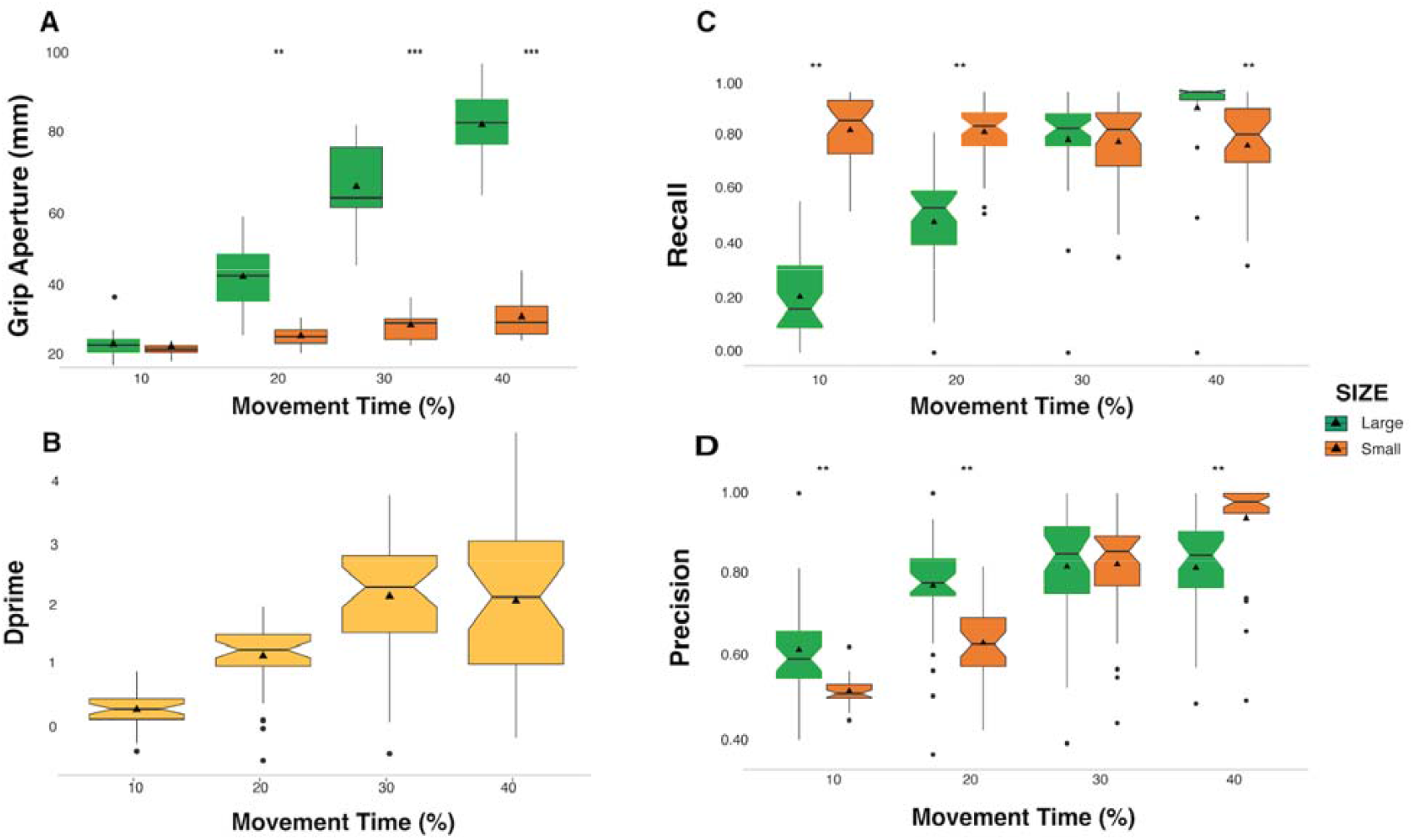
**A**. *Grip Aperture*. Boxplots with individual data points of the 16 selected reach-to-grasp actions for large and small targets. Results highlight significant differences in grip aperture at 20%, 30%, and 40% of the movement. \*\**p* < .01; \*\*\**p* < .001. **B**. *Dprime. d’* increases monotonically from 10% to 30% of the movement time. **C**. *Recall*. Boxplots of participants’ Recall for the 16 selected video clips. Participants had higher Recall of the small target at 10% and 20% and higher Recall for the large target at 40% of the movement. **D**. *Precision*. Participants show higher Precision for the large target at 10% and 20% and higher Precision for the small target at 40% of the movement. \*\**p* < .01

Results on the analysis of grip aperture on the full dataset (160 actions) mirrored the results from the selected dataset (16 actions): we found a main effect of Time and Size, as well as a statistically significant Size*Time interaction (all ps <.001). These results are depicted in Figure S1. Full sanity check results are reported in Table S1A. Post-hoc tests corrected for Bonferroni also reflected previous results, with significantly enhanced grip aperture for Large compared to Small targets at 20%, 30%, and 40% (all ps <.001), but not at 10% (p =.518) of movement time.

#### Wrist Velocity

We found a main effect of Time (F(1.652,9.913)= 338.470, p < .001, pη = 0.983). Conversely, no main effects or interactions involving the factor Size were significant (all ps > .05). Bonferroni corrected post-hoc comparison of the main effect of Time showed a significant increase in wrist velocity between each movement time (all ps < .005) except 30% compared to 40% time of movement (p = 1), (mean±SE; 10%: 271.39±10.39; 20%: 463.84±16.51; 30%: 569.26±17.96; 40%: 583.70±12.66). Results on the analysis of wrist velocity on the full dataset (160 videos) revealed a main effect of Size (p=.048), Time (p <.001), and a Size*Time interaction (p=.007); see Table S1B for full sanity check results. Post-hoc comparisons corrected for Bonferroni revealed significantly enhanced velocity for Large compared to Small targets only at 40% of movement (p=.026, all other ps = 1).

### Target Prediction

*d’*. Analysis on *d’* revealed a main effect of Time (F(1.804, 64.946)=52.136, p<.001, *pη* = 0.592, G-G corrected). Bonferroni corrected post-hoc test confirmed a monotonic increase of *d’* from 10% to 30% (all ps <.001). There was no significant difference between 30% and 40% (p = 1), confirming that target discrimination was already maximal at 30% (mean±SD: 10%: 0.273±0.043; 20%: 1.095±0.084; 30%: 2.005±0.149; 40%: 1.936±0.202), see Figure 2B.

#### Recall

Analysis of Recall on the 2×4 repeated-measures ANOVA showed a main effect of Time (F(3,108)= 177.971, p < .001, pη = 0.832) and Size (F(1,36)= 37.547, p < .001, pη = 0.511) as well as a statistical interaction Time*Size (F(3,108)= 154.543, p < .001, pη = 0.811). This result confirms that our sample size was sufficient to detect the medium effect of interest. Post-hoc analysis with Bonferroni correction on the Time*Size interaction showed greater Recall for the Small target compared to the Large target at 10% and 20% of movement time (mean±SE; 0.844±0.021 vs. 0.215±0.025 at 10%; 0.840±.018 vs. 0.499±0.031 at 20%; p < .001). At 40% of the movement time, the Recall was greater for the Large target (mean±SE; 0.930±0.008 vs 0.784±0.032; p = .001). There was no difference in Recall for the Large and Small targets at 30% of the time video (mean±SE; 0.816±0.030 vs 0.808±0.026, p = .852). These results are illustrated in Figure 2C. Bonferroni corrected post-hoc test on the main effect of Size showed higher Recall for the small targets than the large ones (mean±SE; 0.820±0.20 vs. 0.615±0.023, p = .001). The same test on the main effect of Time highlighted significantly larger Recall for 40% compared to 30%, 20%, and 10%, larger Recall for 30% compared to 20% and 10%, and larger Recall for 20% compared to 10% (mean±SE; 0.859±0.021 vs. 0.812±0.20 vs. 0.670±0.015 vs. 0.529±0.006, all ps < .001)..

#### Precision

Analysis of Precision showed a significant main effect of Time (F(3,105)= 193.683, p < .001, pη^2^ = 0.847 as well as a significant interaction between Time*Size (F(3,105)= 57.566; p < .001; pη^2^ = 0.662). Again, this result confirms that our sample size was sufficient to detect the medium effect of interest. The main effect of Size was not significant (p = .08). Bonferroni corrected post-hoc comparison on the Time*Size interaction highlighted higher Precision for the Large compared to the Small target at 10% (mean±SE; 0.622±0.022 vs. 0.522±0.005) and 20% (mean±SE; 0.776±0.020 vs. 0.642.014). At 40% of the movement time, Precision was higher for Small targets than Large ones (mean±SE; 0.944±0.18 vs 0.819±0.22), all ps < .01. Bonferroni corrected post-hoc test on the main effect of Time yielded significantly larger Precision for 40% compared to 30%, 20%, and 10%; larger Precision for 30% compared to 20% and 10%, and larger Precision for 20% compared to 10% (p < .001; mean± SE; 0.881±.018; 0.829±.019; 0.709±.014; 0.572±.012). These results are illustrated in Figure 2D..

Reaction Time. RT analysis showed a significant main effect of Time (F(3,105) = 98.880, p < .001, pη = .739) and a significant Size*Time interaction (F(3,105) = 29.927, p < .001, pη = 0.461). Post-hoc analysis with Bonferroni correction revealed faster RTs for the Small compared to Large targets at 10% (p = .004; mean±SE; 615.793 ± 30.278 vs. 672.565 ± 31.034) and faster RTs for the Large targets compared to the Small ones at 30% (p < .001; mean±SE; 413.465 ± 15.645 vs. 478.288 ± 18.218) and 40% (p < .001; mean±SE; 371.596 ± 13.760 vs. 457.412±18.432). These results are illustrated in Figure S2.

### Decision Tree

We trained a Decision Tree (DT) to predict participants’ responses at different movement times. At 10%, the DT classified 100% of participants’ responses as “Small,” with an accuracy of 81% (explained by the fact that participants answered “Small” in 81% of trials). The DT analysis included the grip apertures at 0% and 10% of the movement time as predictors. Notably, it discarded information on the velocity of movement.

At 20% of the movement, the DT classified 89% of participants’ responses as “Small” and 52% as “Large” with an accuracy of 76.5%. In this case, the analysis’s independent variables were the grip apertures at 20%, 10%, and 0% of the movement time.

At 30%, the DT classified 89.9% of participants’ responses as “Small” and 75.2% as “Large” with an accuracy of 82.3%. In this case, the analysis’s independent variables were the grip apertures at 30%, 0%, and 20% of the movement time.

Finally, at 40%, the DT classified 94.6% of participants’ responses as “Small” and 81.3% as “Large”, with an overall accuracy of 87%. The selected independent variables, in this case, were the grip apertures at 30% and 0% and velocity at 30%. Nevertheless, the model achieved similar results when selecting only grip aperture as predictors (Small: 94.6%; Large: 81.3%; accuracy: 87%; predictors: grip apertures at 40%, 0%, and 20% of the movement time). A summary of these results is described in Table S2.

### Classification

#### SVM classifier

The overall accuracy for the SVM classifier was 0.70 at 10% of the movement time, 0.87 at 20%, 0.93 at 30%, and 0.95 at 40%. Analyses on *Recall* for each size (Large/Small) revealed that the SVM classifier yielded 0.49 for the large target and 0.91 for the small target at 10%. At 20%, the SVM classifier yielded a Recall of 0.83 and 0.90 for the large and small targets, respectively. At 30% and 40% of the movement time, the classifier showed high Recall both for the large target (30%: 0.88, 40%: 0.92) and the small target (30%: 0.98, 40%: 0.98). Analyses on Precision showed that the SVM classifier yielded 0.74 for the large target and 0.66 for the small target at 10% of the movement time, 0.90 for the large target and 0.86 for the small target at 20%, 0.98 for the large and 0.92 for the small at 30%, 0.98 for the large target and 0.94 for the small target at 40% of the movement time.

#### CNN-RNN classifier

The CNN-RNN model (whose hyperparameters were set as specified in Section 3.4) was tested over video clips displaying 10%, 20%, 30%, and 40% of the movement time. The overall accuracy was slightly lower than that of the SVM classifier (10%: 0.48; 20%: 0.69; 30%: 0.78; 40%: 0.90).

At 10% of the movement time, the neural network yielded a *Recall* of 0.14 for the large target and 0.81 for the small target. At 20%, Recall for the large target was 0.60 and 0.78 for the small target. At 30%, the Recall was 0.71 for the large target and 0.85 for the small target, and at 40%, it was 0.88 for the large target and 0.83 for the small target.

The *Precision* of the CNN-RNN classifier was 0.42 for the large target and 0.49 for the small target at 10%; 0.73 for the large target and 0.55 for the small target at 20%; 0.83 for the large target and 0.75 for the small target at 30%; 0.92 for the large target and 0.88 for the small target at 40%. Results for Precision and Recall in participants, SVM, and CNN-RNN are shown in Figure 3 and summarized in Table S3.

**Figure 3.**
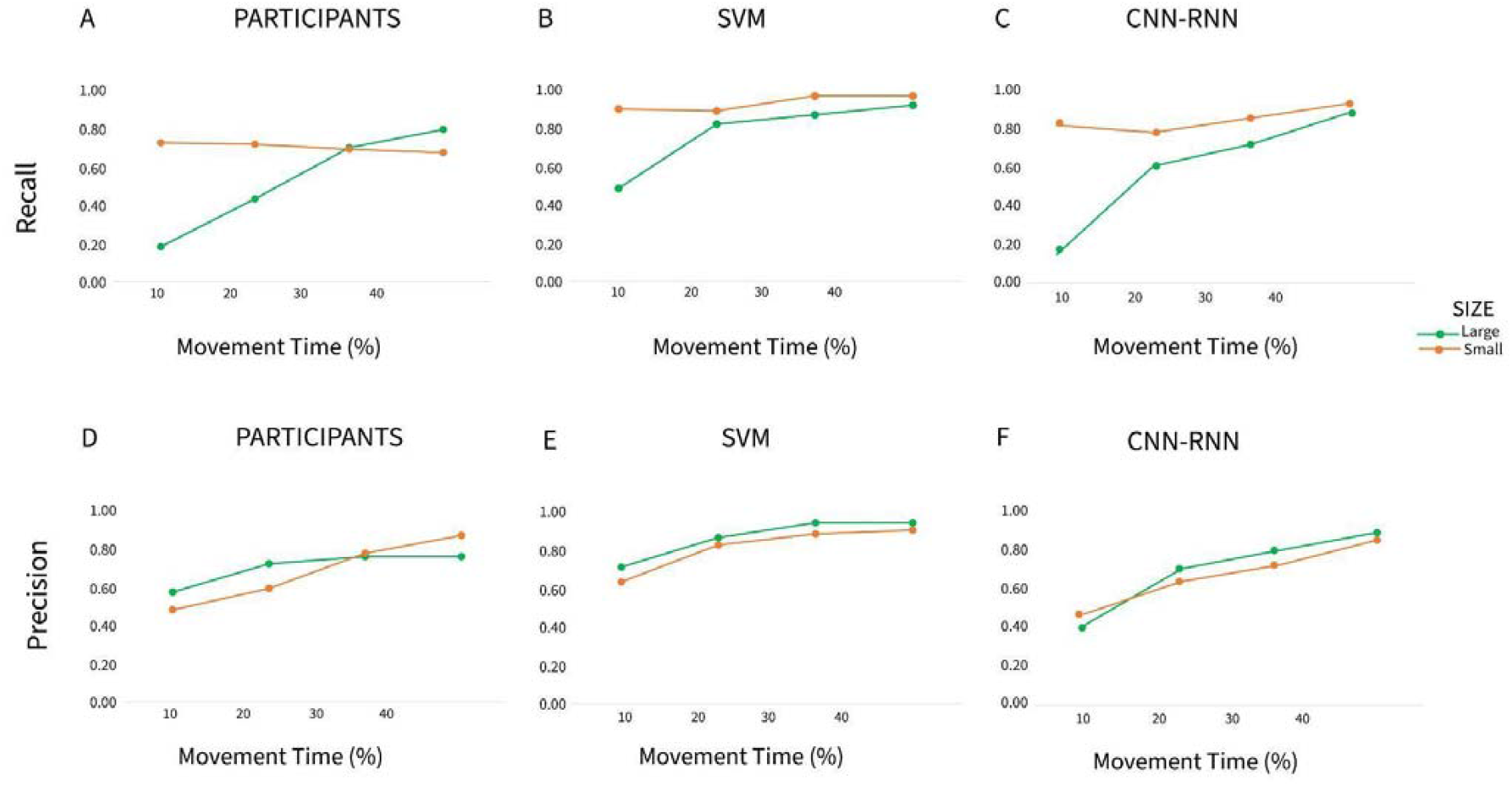
Participants, SVM, and CNN-RNN Recall and Precision. *Upper panel*: **A** Participants, **B** SVM, and **C** CNN-RNN classifier performances on Recall for large and small targets. *Lower panel*: **D** Participants, **E** SVM, and **F** CNN-RNN classifier performances on Precision for large and small targets.

### Agreement between SVM, CNN-RNN, and Participants

To evaluate the agreement between SVMs, CNN-RNNs, and human participants, we use Light Cohen’s Kappa. When comparing SVMs to participants, Light Cohen’s K is 0.17 at 10% (poor agreement), 0.55 at 20% (fair agreement), 0.72 at 30% (good agreement), and 0.78 at 40% (excellent agreemen) of movement time. For CNN-RNNs, Light Cohen’s K is 0.08 at 10% (poor agreement), 0.22 at 20% (poor agreement), 0.51 at 30% (fair agreement), and 0.73 at 40% (good agreement).

We also calculated Light Cohen’s Kappa to assess the agreement between SVMs, CNN-RNNs, and participants for large and small objects separately. When we consider only large targets, Light Cohen’s K for SVM is 0.18 at 10% (poor agreement), 0.38 at 20% (poor agreement), 0.42 at 30% (fair agreement), and 0.89 at 40% (excellent agreement). The agreement between CNN-RNNs and participants for large targets showed Light Cohen’s K of 0.01 at 10%, -0.01 at 20%, 0.02 at 30%, and 0.67 at 40%, indicating poor agreement in all cases except at 40%, where agreement is good. For small objects, Light Cohen’s K for SVM is 0.32 (poor) at 10%, 0.65 at 20% (good), 0.43 at 30% (fair), and 0.46 at 40% (fair). Concerning agreement between CNN-RNNs and participants, Light Cohen’s K is 0.02 at 10%, 0.03 at 20%, 0.20 at 30%, and 0.30 at 40%, consistently indicating poor agreement. These results are summarized in Figure 4 and Table S4.

**Figure 4.**
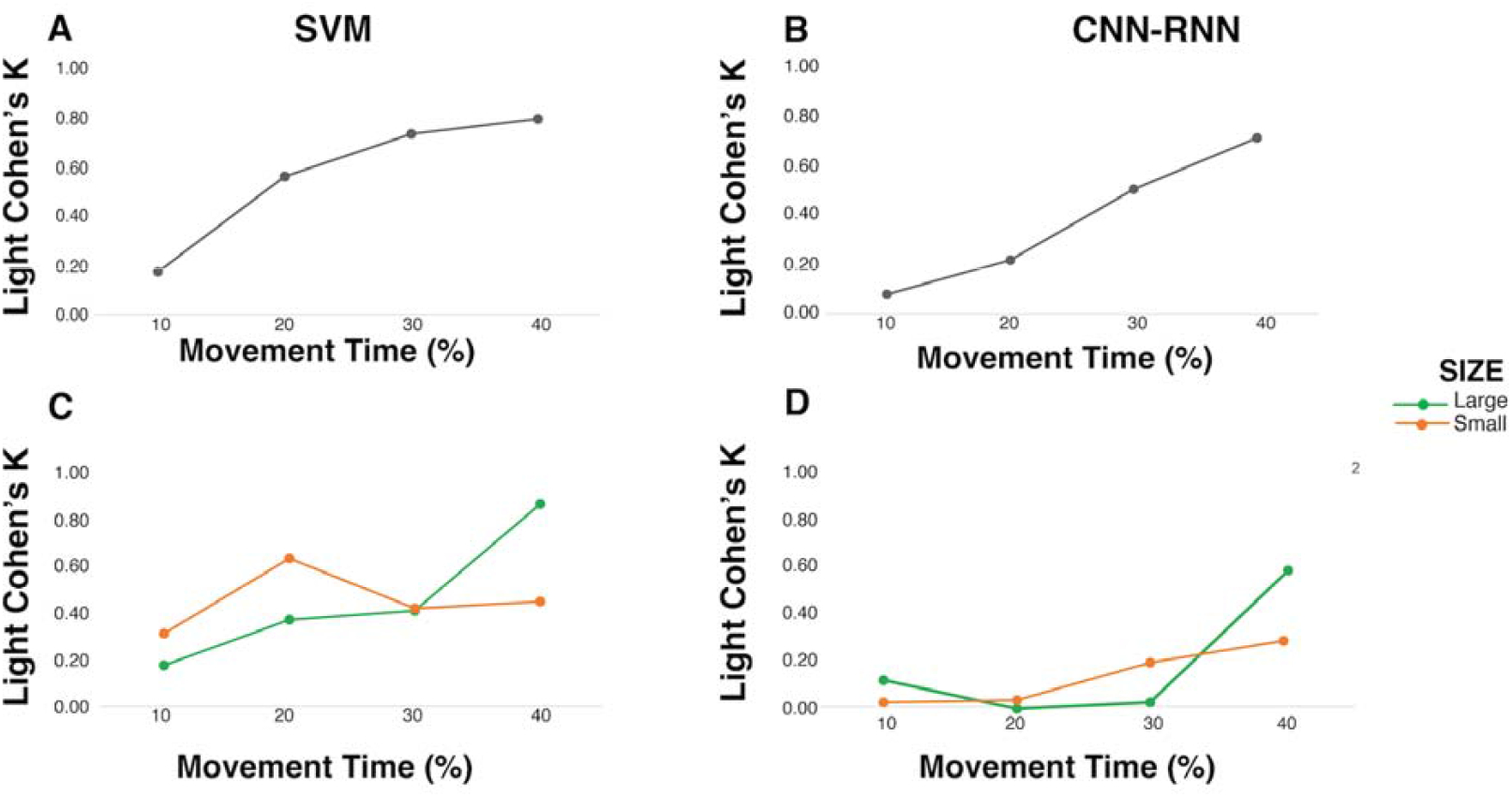
SVM and CNN-RNN agreement with participants’ performance in target prediction task. *Upper panel*: Light Cohen’s Kappa measuring overall agreement of A. SVMs and B. CNN-RNNs with participant performance (grey line). *Lower panel*: Light Cohen’s Kappa measuring agreement of C. SVMs and D. CNN-RNNs with participant performance for large targets (green line) and small targets (orange line). Both panels demonstrate that SVMs showed higher agreement with participant performance than CNN-RNNs.

## DISCUSSION

This study investigated target prediction during early-phase action observation. While prior research established successful target discrimination in early grasping phases (Ansuini et al., 2016), the underlying predictive process remains unexplored. Specifically, how do predictions vary across different target types throughout movement observation? Do these prediction variations relate to differences in information processing, with different cues relevant for different targets at different timepoints? By addressing these questions through a target prediction task, we aimed to uncover the temporal dynamics and cognitive mechanisms underlying action prediction.

Participants should infer the target size (small or large) by observing video clips presenting a schematic (via red-stick diagrams) rendition of the kinematics of grasping actions executed with a precision grip or whole-hand prehension. The video clips showed the initial 10%, 20%, 30%, or 40% of the movement time. After data collection, we systematically analyzed the data with statistical and machine learning methods to unravel the potential strategies employed by human and artificial agents in early target identification.

We first calculated the *d’* for each participant to measure target prediction performance. The results showed that participants discriminated between large and small targets from 20% of the movement time. Moreover, d’ monotonically increased from 10% to 30% but did not significantly differ from 30% to 40%, suggesting that, at 30% of the movement time, participants’ performance in target prediction was already maximal. As in Ansuini et al. (2016), target discrimination performance mirrored the kinematic information, notably grip aperture. Indeed, we found that the grip aperture differentiated between small and large targets from 20% up to 40% of the observed actions. No differences were found at 10%.

*d’* measures the overall detection sensitivity but does not tell us anything about potential target-related differences in prediction strategy. To address this issue, we separately computed Recall (i.e., the amount of correct target identification on the total target presentation) and Precision (i.e., the amount of correct target identification on the total target choice) for each target. Results showed asymmetric prediction outcomes for large and small targets over time. At 10%, when the grip aperture did not provide reliable cues for distinguishing between small and large targets, participants’ responses were strongly biased towards the small target, likely because the observed hand was initially in a pinched/closed position. This bias was evidenced by Recall scores for the small target four times higher than the large target and significantly faster RTs for the small target than the large target.

From 20% to 40%, as kinematic information increasingly allowed for discrimination between small and large targets, we observed a progressive increase in participants’ Recall for large targets while their Recall for small targets remained constant. Participants likely relied on maximal grip aperture as the primary cue for target size prediction. Kinematic analysis of grasping actions presented as stimuli showed that the maximum grip aperture discriminated between the types of grip required by the two different targets from 20% up to 40% of action, while this did not apply to wrist velocity. Since the grasping actions began with a pinched/closed hand, participants defaulted to small target predictions until grip aperture signaled full-hand grasping. This created a systematic bias toward small targets during the very early movement phase when kinematic information was sparse, but large target recall improved as more information became available throughout the movement.

A differential pattern was also found in participants’ Precision. Similar to Recall, Precision increased over time but not uniformly. At 10% and 20%, Precision was higher for large targets than for small ones. However, by 40%, this pattern reversed, with Precision being higher for small targets than for large ones.

We employed a Decision Tree (DT) algorithm to model participants’ prediction strategy using the maximum grip aperture and wrist velocity at each movement time as predictors. The results revealed a pattern consistent with participants’ asymmetric prediction outcomes for small and large target prediction over time. At 10%, the DT classified the participants’ responses as “Small” 100% of the time. At 20%, 30%, and 40%, the decision tree progressively increased the classification of participants’ responses as “Large” (52%, 75.2%, and 81.3%, respectively), while the classification of participants’ responses as “Small” remained nearly constant (around 89%–94.6%).

To better understand the dynamics of the prediction strategy over time, we investigated the performance in target discrimination of two machine learning algorithms and compared them to participants’ performance. The first algorithm we employed was a Support Vector Machine (SVM). We trained this algorithm on the kinematic parameters previously extracted from each movement time of the executed reach-to-grasp actions. The second algorithm was a Neural Network with a CNN-RNN architecture, which was trained directly on the portions of video clips corresponding to the reach-to-grasp actions at each movement time. Overall, both algorithms improved their classification accuracy over time. As expected, the SVM became highly accurate much earlier than the CNN-RNN architecture (0.70 at 10% and 0.87 at 20% vs 0.48 at 10% and 0.69 at 20%). As for Recall and Precision, both algorithms aligned with the participants’ response trends: they exhibited a progressive increase in the Recall rate for the large target, with the Recall rate for the small target remaining constant, and an overall increase in the Precision rate for both targets. However, some differences exist: SVM generally performed better in the early stages. In particular, there were higher Recall values for the “Large” responses and higher Precision values for the “Small” responses at 10% and 20%.

We computed Light Cohen’s K on a single-trial basis to quantify the degree of agreement between participants, SVM, and CNN-RNN responses. According to Light Cohen’s Kappa, SVMs outperformed CNN-RNNs, meaning SVMs matched participants’ predictions more than CNN-RNNs. Note that, at 10%, both algorithms showed poor agreement with participants. At 30% and 40%, SVMs exhibited good and excellent agreement with participants’ predictions, respectively, while CNN-RNNs reached a good agreement at 40% only.

We also computed Light Cohen’s K on a single-trial basis to quantify the agreement between SVMs, CNN-RNNs, and participants’ prediction outcomes for large and small targets separately. SVMs agreed with participants at certain movement times, even if the trend was not uniform between targets. With large targets, the agreement increased monotonically, reaching its peak at 40% (excellent agreement). With the small targets, the agreement became rapidly good (at 20%), but it decreased to fair at 30% and 40%. Unlike SVMs, CNN-RNNs exhibited a very poor agreement for both targets, except for the large target at 40% (good agreement).

Our results suggest that kinematic information influences participants’ performance asymmetrically across target types over time. This asymmetry appears in both behavioral data and decision tree predictions based on kinematic parameters: as more kinematic information becomes available, large target predictions increase while small target predictions remain constant. Similar asymmetric patterns emerge in the SVM and CNN-RNN algorithms, confirming this fundamental prediction strategy.

Comparing these algorithms offers insights into the underlying processes of participants’ prediction performance. The SVM processed kinematic parameters relevant to action execution and control (Ansuini et al., 2015; Khera & Kumar, 2020), while the CNN-RNN extracted spatiotemporal patterns from video frames of grasping actions, simulating higher-order visual processing (Karpathy et al., 2015). As expected, the SVM outperformed the CNN-RNN during early processing stages. Notably, the SVM progressively aligned with participants’ responses, particularly for large target predictions, a pattern not observed with the CNN-RNN.

Previous research supports the idea that processing the kinematic parameters relevant to action execution and control significantly impacts participants’ performance in target prediction tasks. Costantini et al. (2014) asked participants to view videos of a hand grasping a small or large tomato without providing them with prior information about the potential targets. In a control condition, participants had to watch videos of the hand reaching for one of the two targets with a closed fist without pre-shaping any grip. Participants demonstrated significantly faster and more accurate proactive gaze towards the targeted tomato in the experimental condition compared to the control (see also Ambrosini et al., 2011). Notably, when repetitive transcranial magnetic stimulation (rTMS) was applied to the left ventral premotor cortex (PMv), participants’ gaze proactivity drastically decreased, suggesting an impaired ability to use hand pre-shaping information. In contrast, rTMS application to a high-order visual cortical area, such as the posterior portion of the superior temporal sulcus, produced no comparable effects.

This aligns with numerous studies demonstrating that observing hand-grasping actions activates the same motor brain regions engaged when the observer executes and controls the actions themselves. The motor cortical activation facilitates the transformation of the observed action’s sensory representation into the observer’s motor representation, enabling immediate comprehension of the action’s goal and target (Rizzolatti et al., 2001; Rizzolatti & Sinigaglia, 2010, 2016). A multivariate fMRI study has demonstrated that action goals are mainly coded in the inferior parietal cortex, while several parietal and premotor structures process the target-related grip parameters (Errante et al., 2021). Notably, this process occurs whether the action target is directly observed or inferred (Villiger et al., 2011). When the target must be inferred, the observer typically engages the parietal and premotor cortices to a greater extent than when the target is visible. In contrast, visible target encoding primarily relies on areas within the ventral visual pathway (Thiaux & Keysers, 2015).

While our findings converge with evidence that action observation engages motor resources, we make three key advances beyond prior research. First, we demonstrate that processing of kinematic information supports target-specific prediction performance, with asymmetric predictive outcomes over time. Second, our machine learning comparisons provide computational evidence that this target-specific performance emerges from processing kinematic features relevant to action execution and control rather than visual patterns of the observed action. Finally, by tracking prediction strategies from the earliest observation phases, we reveal how target-dependent kinematic features shape action understanding from movement onset. These findings have significant implications for robot-human interactions in manufacturing and neuro-rehabilitation settings, specifically enabling robots to utilize and manipulate cues for anticipating movement targets (Boucher et al., 2012).

A potential limitation of our study is that the observed target-related asymmetry in predictive outcomes depends on the initial hand posture of the observed action. However, our chosen starting posture aligns with standard kinematic studies, and notably, no prior action observation research has examined the prediction strategy. Moreover, starting with a fully open rather than pinched/closed hand would likely yield an opposite asymmetric pattern. Finally, the finding that action prediction depends on initial hand posture and varies by target type reflects real-world scenarios, where actions are observed under varying conditions that require modification of predictive strategy.

A further limitation is our focus on predicting outcomes without examining participants’ subjective confidence in their predictions. Future research should investigate whether prediction confidence increases uniformly across time and targets or shows target-specific patterns that mirror the observed prediction outcome asymmetries. Understanding how prediction accuracy and subjective confidence relate could reveal how different cognitive components integrate during action prediction.

## ACKNOWLEDGMENTS

This article was supported by the Department of Philosophy ‘Piero Martinetti’ of the University of Milan with the Project “Departments of Excellence 2018-2022” awarded by the Italian Ministry of Education, University and Research (MIUR) (to CS), and by the PRIN 2017 project “The cognitive neuroscience of interpersonal coordination and cooperation: a motor approach in humans and non-human primates” (201794KEER; to CS).

## AUTHOR CONTRIBUTIONS

Conceptualization, M.F., M.R., M.Fe, and C.S..; Methodology, M.F., F.D., G.B., and D.Q.; Software, F.D., and D.Q; Formal analysis, M.F., F.D., and G.B.0; Investigation, M.F., and D.Q.; Writing –Original Draft, C.S., M.F., F.D., and G.B.; Writing – Review and Editing, C.S., M.F., F.D., G.B., M.R., and M.Fe..; Visualization, M.F. and F.D.; Supervision, C.S., M.R., M.Fe.; Funding Acquisition C.S.

## ADDITIONAL INFORMATION

Supplemental Tables and Figures can be found in ***Supplemental Materials***.

## References

1. Ambrosini, E., Costantini, M., & Sinigaglia, C. (2011). Grasping with the eyes. Journal of Neurophysiology, 106(3), 1437–1442.

2. Ambrosini, E., Pezzulo, G., & Costantini, M. (2015). The eye in hand: Predicting others’ behavior by integrating multiple sources of information. Journal of Neurophysiology, 113(7), 2271–2279.

3. Ambrosini, E., Reddy, V., De Looper, A., Costantini, M., Lopez, B., & Sinigaglia, C. (2013). Looking ahead: Anticipatory gaze and motor ability in infancy. PloS One, 8(7), e67916.

4. Ambrosini, E., Sinigaglia, C., & Costantini, M. (2012). Tie my hands, tie my eyes. Journal of Experimental Psychology: Human Perception and Performance, 38(2), 263.

5. Ansuini, C., Cavallo, A., Koul, A., D’Ausilio, A., Taverna, L., & Becchio, C. (2016). Grasping others’ movements: Rapid discrimination of object size from observed hand movements. Journal of Experimental Psychology: Human Perception and Performance, 42(7), 918.

6. Ansuini, C., Cavallo, A., Koul, A., Jacono, M., Yang, Y., & Becchio, C. (2015). Predicting Object Size from Hand Kinematics: A Temporal Perspective. PLoS ONE 10(3): e0120432.

7. Becchio, C., Cavallo, A., Begliomini, C., Sartori, L., Feltrin, G., & Castiello, U. (2012). Social grasping: From mirroring to mentalizing. Neuroimage, 61(1), 240–248.

8. Campbell, J. I. D., & Thompson, V. A. (2012). MorePower 6.0 for ANOVA with relational confidence intervals and Bayesian analysis. Behavior Research Methods, 44(4), 1255–1265. 10.3758/s13428-012-0186-0

9. Carpinella, I., Mazzoleni, P., Rabuffetti, M., Thorsen, R. & Ferrarin, M. (2006). Experimental protocol for the kinematic analysis of the hand: definition and repeatability. Gait & Posture 23, 445–454.

10. Castiello, U. (2005). The neuroscience of grasping. Nature Reviews Neuroscience, 6(9), 726–736.

11. Cohen, J. (1960). A coefficient of agreement for nominal scales. Educational and Psychological Measurement, 20(1), 37–46.

12. Costantini, M., Ambrosini, E., & Sinigaglia, C. (2012a). Out of your hand’s reach, out of my eyes’ reach. Quarterly Journal of Experimental Psychology, 65(5), 848–855.

13. Costantini, M., Ambrosini, E., & Sinigaglia, C. (2012b). Does how I look at what you’re doing depend on what I’m doing? Acta Psychologica, 141(2), 199–204.

14. Costantini, M., Ambrosini, E., Cardellicchio, P., & Sinigaglia, C. (2014). How your hand drives my eyes. Social Cognitive and Affective Neuroscience, 9(5), 705–711.

15. Cristianini, N., & Shawe-Taylor, J. (2000). An Introduction to Support Vector Machines and Other Kernel-based Learning Methods. (Cambridge University Press).

16. Errante, A., Ziccarelli, S., Mingolla, G. P., & Fogassi, L. (2021). Decoding grip type and action goal during the observation of reaching-grasping actions: A multivariate fMRI study. NeuroImage, 243, 118511.

17. Fleiss, J. L., Levin, B. & Paik, M. C. (2003). Statistical Methods for Rates and Proportions (John Whiley & Sons).

18. Jakobson, L. S., & Goodale, M. A. (1991). Factors affecting higher-order movement planning: a kinematic analysis of human prehension. Experimental Brain Research, 86, 199–208.

19. Jeannerod M. (1981). Intersegmental coordination during reaching at natural visual objects, in Attention and Performance, Vol. IX, eds Long J., Baddeley A. (Hillsdale, NJ: Erlbaum), 153–168.

20. Jeannerod, M. (1984). The timing of natural prehension movements. Journal of Motor Behavior, 16(3), 235–254.

21. Jeannerod, M., Arbib, M. A., Rizzolatti, G., & Sakata, H. (1995). Grasping objects: the cortical mechanisms of visuomotor transformation. Trends in neurosciences, 18(7), 314–320.

22. Kapoor, A., Gulli, A., & Pal, S. (2023). Deep Learning with TensorFlow and Keras–3rd edition. Small, 362, 362.

23. Karpathy, A., Johnson, J., & Fei-Fei, L. (2015). Visualizing and understanding recurrent networks. arXiv preprint 1506.02078

24. Khera, P., & Kumar, N. (2020). Role of machine learning in gait analysis: A review. Journal of Medical Engineering & Technology, 44(8), 441–467.

25. Krishnan-Barman, S., Forbes, P. A., & Hamilton, A. F. D. C. (2017). How can the study of action kinematics inform our understanding of human social interaction?. Neuropsychologia, 105, 101–110.

26. Light, R. J. (1971). Measures of response agreement for qualitative data: some generalizations and alternatives. Psychological Bulletin, 76(5), 365.

27. Lindsay G. W. (2021). Convolutional Neural Networks as a Model of the Visual System: Past, Present, and Future. Journal of Cognitive Neuroscience, 33(10), 2017–2031.

28. Lipton, Z. C., Berkowitz, J., & Elkan, C. (2015). A critical review of recurrent neural networks for sequence learning. arXiv preprint 1506.00019.

29. Pedregosa, F., Varoquaux, G., Gramfort, A., Michel, V., Thirion, B., Grisel, O., Blondel, M., Prettenhofer, P., Weiss, R., Dubourg, V., Vanderplas, J., Passos, A., & Cournapeau, D. (2011). Scikit-learn: Machine Learning in Python. Journal of Machine Learning Research, 12, 2825–2830.

30. Paulignan, Y., Frak, V. G., Toni, I., & Jeannerod, M. (1997). Influence of object position and size on human prehension movements. Experimental Brain Research, 114, 226–234.

31. Paulignan, Y., Jeannerod, M., MacKenzie, C., & Marteniuk, R. (1991). Selective perturbation of visual input during prehension movements: 2. The effects of changing object size. Experimental Brain Research, 87, 407–420.

32. Piedimonte, A., Garbarini, F., Rabuffetti, M., Pia, L., Montesano, A., Ferrarin, M., & Berti, A. (2015). Invisible grasps: Grip interference in anosognosia for hemiplegia. Neuropsychology, 29(5), 776–781.

33. Rizzolatti, G., & Sinigaglia, C. (2010). The functional role of the parieto-frontal mirror circuit: interpretations and misinterpretations. Nature Reviews Neuroscience, 11(4), 264–274.

34. Rizzolatti, G., & Sinigaglia, C. (2016). The mirror mechanism: a basic principle of brain function. Nature Reviews Neuroscience, 17(12), 757–765.

35. Rizzolatti, G., Fogassi, L., & Gallese, V. (2001). Neurophysiological mechanisms underlying the understanding and imitation of action. Nature Reviews Neuroscience, 2(9), 661–670.

36. Thioux, M., & Keysers, C. (2015). Object visibility alters the relative contribution of ventral visual stream and mirror neuron system to goal anticipation during action observation. NeuroImage, 105, 380–394.

37. Villiger, M., Chandrasekharan, S., & Welsh, T. N. (2011). Activity of human motor system during action observation is modulated by object presence. Experimental Brain Research, 209, 85–93.

38. Whitwell, R. L., & Goodale, M. A. (2013). Grasping without vision: time normalizing grip aperture profiles yields spurious grip scaling to target size. Neuropsychologia, 51(10), 1878–1887.

