## Supplemental Material for "Early Target Prediction in Action Observation"

**SUPPLEMENTAL MATERIALS**

**Table S1**

Results of the 2*2*2*2*4 Mixed Repeated-Measures ANOVA (between-factor: Agent (Female, Male); within-factors Side (Right, Left), Distance (60, 70), Size (Large, Small), Time (10%, 20%, 30%, 40%)) for the whole dataset of grasping movements (160 videos) on Grip Aperture (A) and Wrist Velocity (B).

*p <.05; *p <.01

| A. **Grip Aperture** | | | | |
| --- | --- | --- | --- | --- |
|  | **df** | **F** | **p** | **η²_p_** |
| Side | 1 | 11.535 | 0.003** | 0.391 |
| Side ✻ Agent | 1 | 7.493 | 0.014* | 0.294 |
|  | 18 |  |  |  |
| Distance | 1 | 0.411 | 0.530 | 0.022 |
| Distance ✻ Agent | 1 | 7.187 | 0.015 | 0.285 |
|  | 18 |  |  |  |
| Size | 1 | 475.185 | <.001** | 0.964 |
| Size ✻ Agent | 1 | 8.941 | 0.008** | 0.332 |
|  | 18 |  |  |  |
| Time | 3 | 863.518 | < .001** | 0.980 |
| Time ✻ Agent | 3 | 6.919 | < .001** | 0.278 |
|  | 54 |  |  |  |
| Side ✻ Distance | 1 | 0.038 | 0.847 | 0.002 |
| Side ✻ Distance ✻ Agent | 1 | 0.619 | 0.442 | 0.033 |
|  | 18 |  |  |  |
| Side ✻ Size | 1 | 54.598 | < .001** | 0.752 |
| Side ✻ Size ✻ Agent | 1 | 0.056 | 0.815 | 0.003 |
|  | 18 |  |  |  |
| Distance ✻ Size | 1 | 0.009 | 0.927 | 4.763×10^-4^ |
| Distance ✻ Size ✻ Agent | 1 | 2.005 | 0.174 | 0.100 |
|  | 18 |  |  |  |
| Side ✻ Time | 3 | 1.315 | 0.279 | 0.068 |
| Side ✻ Time ✻ Agent | 3 | 0.970 | 0.414 | 0.051 |
|  | 54 |  |  |  |
| Distance ✻ Time | 3 | 0.142 | 0.934 | 0.008 |
| Distance ✻ Time ✻ Agent | 3 | 10.500 | < .001** | 0.368 |
|  | 54 |  |  |  |
| Size ✻ Time | 3 | 679.244 | < .001** | 0.974 |
| Size ✻ Time ✻ Agent | 3 | 15.447 | < .001** | 0.462 |
|  | 54 |  |  |  |
| Side ✻ Distance ✻ Size | 1 | 0.426 | 0.522 | 0.023 |
| Side ✻ Distance ✻ Size ✻ Agent | 1 | 0.245 | 0.627 | 0.013 |
|  | 18 |  |  |  |
| Side ✻ Distance ✻ Time | 3 | 0.276 | 0.842 | 0.015 |
| Side ✻ Distance ✻ Time ✻ Agent | 3 | 2.266 | 0.091 | 0.112 |
|  | 54 |  |  |  |
| Side ✻ Size ✻ Time | 3 | 15.887 | < .001** | 0.469 |
| Side ✻ Size ✻ Time ✻ Agent | 3 | 1.388 | 0.256 | 0.072 |
|  | 54 |  |  |  |
| Distance ✻ Size ✻ Time | 3 | 0.478 | 0.699 | 0.026 |
| Distance ✻ Size ✻ Time ✻ Agent | 3 | 2.798 | 0.049 | 0.135 |
|  | 54 |  |  |  |
| Side ✻ Distance ✻ Size ✻ Time | 3 | 2.062 | 0.116 | 0.103 |
| Side ✻ Distance ✻ Size ✻ Time ✻ Agent | 3 | 2.094 | 0.112 | 0.104 |
|  | 54 |  |  |  |
| Agent | 1 | 0.031 | 0.863 | 0.002 |
|  | 18 |  |  |  |
| B. **Wrist Velocity** | | | | |
|  | **df** | **F** | **p** | **η²_p_** |
| Side | 1 | 4.409×10-4 | 0.983 | 2.449×10-5 |
| Side ✻ Agent | 1 | 18.127 | < .001** | 0.502 |
|  | 18 |  |  |  |
| Distance | 1 | 22.621 | < .001** | 0.557 |
| Distance ✻ Agent | 1 | 17.739 | < .001** | 0.496 |
|  | 18 |  |  |  |
| Size | 1 | 4.507 | 0.048* | 0.200 |
| Size ✻ Agent | 1 | 4.515 | 0.048* | 0.201 |
|  | 18 |  |  |  |
| Time | 3 | 1559.081 | < .001** | 0.989 |
| Time ✻ Agent | 3 | 8.981 | < .001** | 0.333 |
|  | 54 |  |  |  |
| Side ✻ Distance | 1 | 3.546 | 0.076 | 0.165 |
| Side ✻ Distance ✻ Agent | 1 | 14.781 | 0.001 | 0.451 |
|  | 18 |  |  |  |
| Side ✻ Size | 1 | 0.032 | 0.860 | 0.002 |
| Side ✻ Size ✻ Agent | 1 | 0.743 | 0.400 | 0.040 |
|  | 18 |  |  |  |
| Distance ✻ Size | 1 | 9.511 | 0.006 | 0.346 |
| Distance ✻ Size ✻ Agent | 1 | 1.600 | 0.222 | 0.082 |
|  | 18 |  |  |  |
| Side ✻ Time | 3 | 0.299 | 0.826 | 0.016 |
| Side ✻ Time ✻ Agent | 3 | 4.703 | 0.005** | 0.207 |
|  | 54 |  |  |  |
| Distance ✻ Time | 3 | 9.623 | < .001** | 0.348 |
| Distance ✻ Time ✻ Agent | 3 | 7.239 | < .001** | 0.287 |
|  | 54 |  |  |  |
| Size ✻ Time | 3 | 4.446 | 0.007** | 0.198 |
| Size ✻ Time ✻ Agent | 3 | 6.114 | 0.001** | 0.254 |
|  | 54 |  |  |  |
| Side ✻ Distance ✻ Size | 1 | 1.563 | 0.227 | 0.080 |
| Side ✻ Distance ✻ Size ✻ Agent | 1 | 0.087 | 0.772 | 0.005 |
|  | 18 |  |  |  |
| Side ✻ Distance ✻ Time | 3 | 11.666 | < .001** | 0.393 |
| Side ✻ Distance ✻ Time ✻ Agent | 3 | 3.043 | 0.037* | 0.145 |
|  | 54 |  |  |  |
| Side ✻ Size ✻ Time | 3 | 3.663 | 0.018 | 0.169 |
| Side ✻ Size ✻ Time ✻ Agent | 3 | 0.142 | 0.934 | 0.008 |
|  | 54 |  |  |  |
| Distance ✻ Size ✻ Time | 3 | 3.303 | 0.027 | 0.155 |
| Distance ✻ Size ✻ Time ✻ Agent | 3 | 1.936 | 0.135 | 0.097 |
|  | 54 |  |  |  |
| Side ✻ Distance ✻ Size ✻ Time | 3 | 1.218 | 0.312 | 0.063 |
| Side ✻ Distance ✻ Size ✻ Time ✻ Agent | 3 | 1.045 | 0.380 | 0.055 |
|  | 54 |  |  |  |
| Agent | 1 | 12.345 | 0.002** | 0.407 |
|  | 18 |  |  |  |

**Table S2**

Results: Decision Tree. Correct classification for participants’ Large and Small responses and overall classification accuracy at 10%, 20%, 30%, and 40%.

|  | 10% | 20% | 30% | 40% |
| --- | --- | --- | --- | --- |
| Correct responses |  |  |  |  |
| *Large* | 0% | 52% | 75.2% | 81.3% |
| *Small* | 100% | 89% | 89.9% | 94.6% |
| *Accuracy* | 81% | 76.5% | 82.3% | 87% |
| Predictors | a10, a0 | a20, a0, a10 | a30, a0, a20 | a40, v30, a20 |

**Table S3**

Results for A. Recall and B. Precision in Human participants, SVM, and CNN-RNN classifiers for large and small targets at 10%, 20%, 30% and 40% of movement time.

| A. **Recall** |  |  |  |  |
| --- | --- | --- | --- | --- |
|  | 10% | 20% | 30% | 40% |
| Human |  |  |  |  |
| *large* | 0.22 | 0.50 | 0.82 | 0.93 |
| *small* | 0.84 | 0.83 | 0.80 | 0.78 |
| SVM |  |  |  |  |
| *large* | 0.49 | 0.83 | 0.88 | 0.93 |
| *small* | 0.91 | 0.9 | 0.98 | 0.98 |
| CNN-RNN |  |  |  |  |
| *large* | 0.14 | 0.60 | 0.71 | 0.88 |
| *small* | 0.81 | 0.78 | 0.85 | 0.83 |

| B. **Precision** |  |  |  |  |
| --- | --- | --- | --- | --- |
|  | 10% | 20% | 30% | 40% |
| Human |  |  |  |  |
| *large* | 0.62 | 0.78 | 0.82 | 0.82 |
| *small* | 0.52 | 0.64 | 0.94 | 0.84 |
| SVM |  |  |  |  |
| *large* | 0.74 | 0.90 | 0.98 | 0.98 |
| *small* | 0.66 | 0.86 | 0.94 | 0.92 |
| CNN-RNN |  |  |  |  |
| *large* | 0.42 | 0.73 | 0.92 | 0.83 |
| *small* | 0.49 | 0.66 | 0.88 | 0.75 |

**Table S4.**

Light Cohen’s Kappa scores (±1*.*96 times standard error) and their interpretation according to Fleiss et al., 2013. A. Overall Light Cohen’s K, for both SVMs and CNN-RNNs classifiers, when compared to participants’ raters B. Light Cohen’s K, showing agreement between participants’ discrimination restricted to the large target, for both SVMs and CNN-RNNs when compared to participants’ raters. C. Light Cohen’s K shows agreement between human participants’ discrimination restricted to the small target for both SVMs and CNN-RNNs when compared to participants’ raters. Notably, SVM has the best agreement according to these metrics.

| A. **Overall** |  |  |  |
| --- | --- | --- | --- |
| 10% | 20% | 30% | 40% |
| SVM |  |  |  |
| 0*.*17±0*.*02 | 0*.*55±0*.*02 | 0*.*72±0*.*02 | 0*.*78±0*.*02 |
| poor | fair | good | excellent |
| CNN-RNN |  |  |  |
| 0*.*08±0*.*01 | 0*.*22±0*.*01 | 0*.*51±0*.*02 | 0*.*73±0*.*02 |
| poor | poor | fair | good |

| B. **Large** |  |  |  |
| --- | --- | --- | --- |
| 10% | 20% | 30% | 40% |
| SVM |  |  |  |
| 0*.*18±0.02 | 0*.*38±0.02 | 0*.*42±0.03 | 0*.*89±0.02 |
| poor | poor | fair | excellent |
| CNN-RNN |  |  |  |
| 0*.*01±0.01 | -0.01±0.01 | 0*.*02±9.01 | 0*.*67±0.02 |
| poor | poor | poor | good |

| C. **Small** |  |  |  |
| --- | --- | --- | --- |
| 10% | 20% | 30% | 40% |
| SVM |  |  |  |
| 0*.*32±0.02 | 0*.*65±0.03 | 0*.*43±0.03 | 0*.*46±0.03 |
| poor | good | fair | fair |
| CNN-RNN |  |  |  |
| 0*.*02±0.01 | 0*.*03±0.02 | 0*.*2±0.02 | 0*.*3±0.02 |
| poor | poor | poor | poor |

**Figure S1**

Grip aperture for Large and Small targets at 10%, 20%, 30%, and 40% of movement in the whole dataset of videos (n = 160). *p <.05; **p <.01

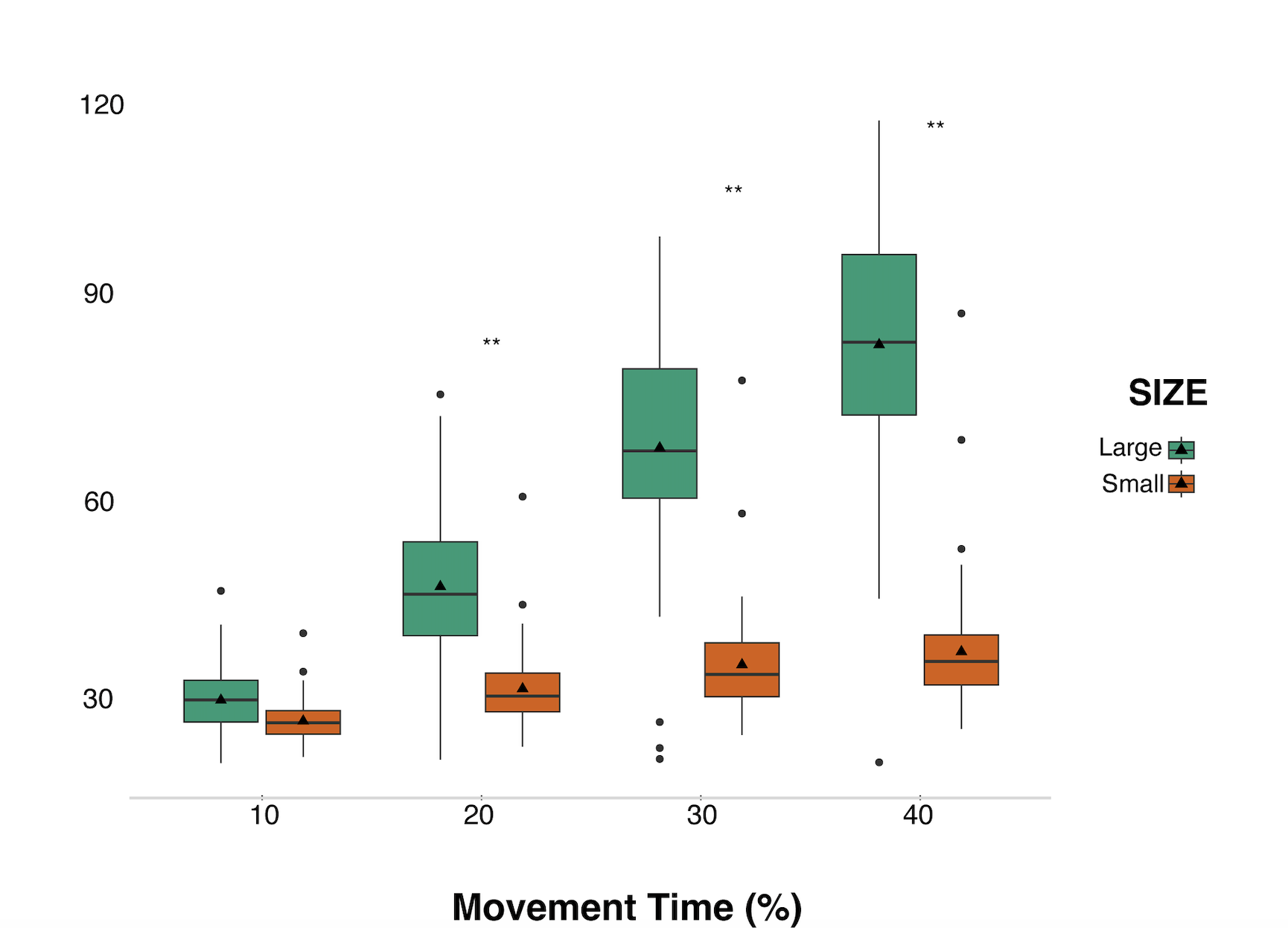

**Figure S2**

Reaction Times (RT) for Large and Small correct responses at 10%, 20%, 30% and 40% of movement time. *p <.05; *p <.01

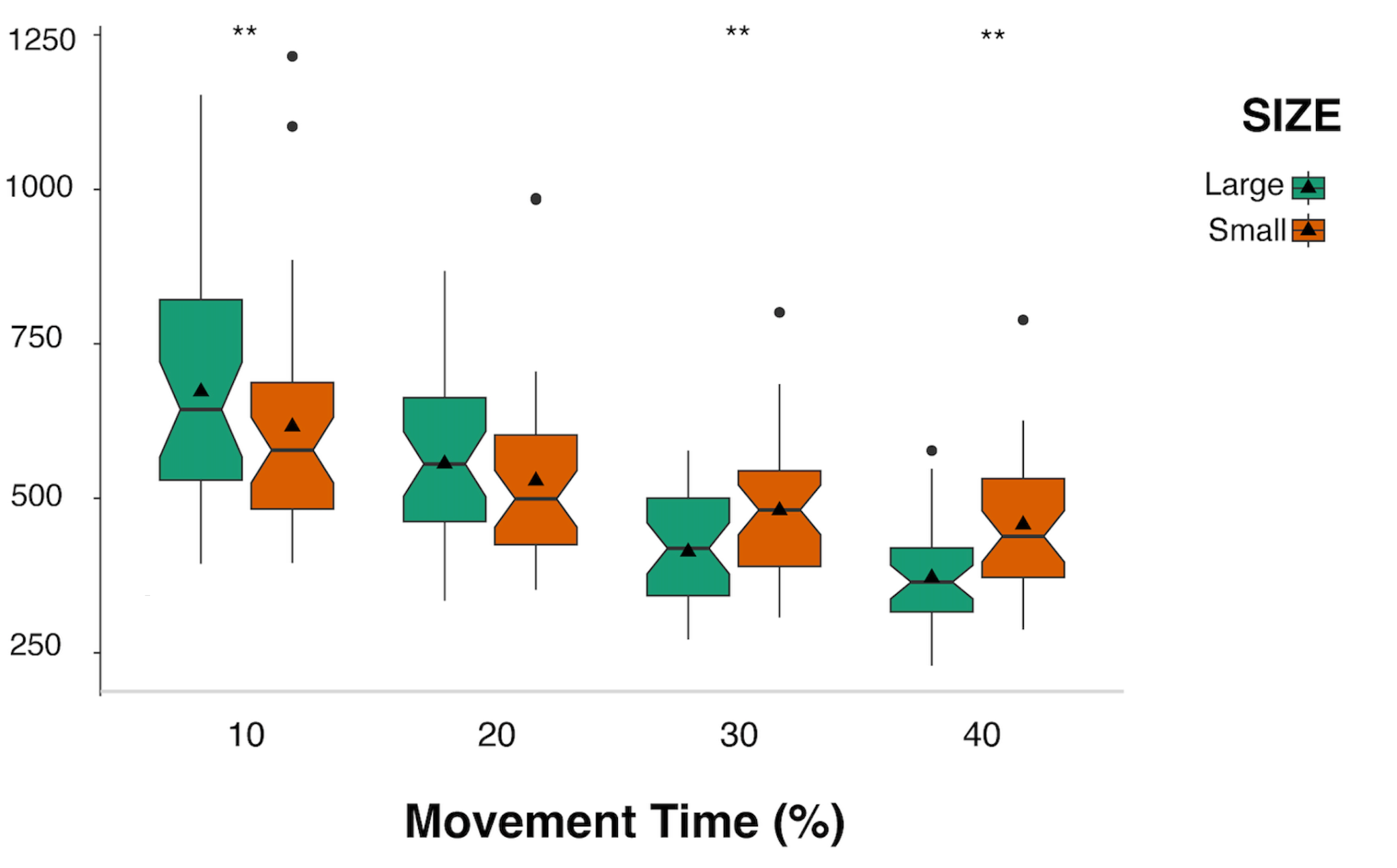
